# An Integrated Computational–Experimental Strategy For the Prediction of Small Molecules as GLP-1R Agonists

**DOI:** 10.64898/2026.03.30.715288

**Authors:** Elena Murcia García, Ning Tian, Jorge-Ricardo Alonso-Fernández, Xiaoqing Cai, Dehua Yang, Juan José Hernández Morante, Horacio Pérez Sánchez

## Abstract

The glucagon-like peptide-1 receptor (GLP-1R) plays a central role in metabolic regulation and is a major therapeutic target for obesity and diabetes. Peptide agonists, like semaglutide, targeting the GLP-1R remain among the most effective regulators of glucose metabolism and appetite. Nonetheless, recent reports about weight regain have limited the effectiveness of GLP1R peptide agonists, sustaining the interest in expanding the chemical diversity of GLP-1R ligands through drug discovery strategies. However, the structural complexity and conformational plasticity of class B1 GPCRs make conventional single-method virtual screening approaches prone to bias and limited chemotype recovery. Using an integrated ligand– and structure-based virtual screening pipeline, explicitly combining complementary ligand-based descriptors, multi-fingerprint similarity, electrostatic similarity, pharmacophore modeling, and multi-conformation docking under a consensus-driven selection strategy, we were able to identify three chemically distinct classes of GLP-1R agonist candidates: GQB47810, a non-peptidic molecule; neuromedin C, a peptide, and 2,5-Pen-enkephalin (DPDPE), a small peptide. From all of them, DPDPE showed the greatest effectiveness, reaching values similar to those of GLP-1, although with lower potency. Further in vitro characterization confirmed that pen-enkephalin behaved as a full agonist and exhibited dual GLP-1R/GIPR agonistic activity.

These findings establish a consensus-driven and transferable computational framework for chemotype-diverse agonist discovery at conformationally flexible GPCR targets, and revealed a pentapeptide with GLP-1-like efficacy as a promising lead for next-generation small peptide therapeutics.

## 1. Introduction

Class B1 G-protein-coupled receptors (GPCRs) are activated by peptide hormones and neuropeptides that regulate essential physiological functions, including neuroprotection, bone turnover, cardiovascular tone, glycemic control, appetite, and energy balance (Bortolato et al., 2014). These receptors predominantly couple to the stimulatory protein G_s_, leading to cyclic adenosine monophosphate (cAMP) production (de Graaf et al., 2016), but they can also engage alternative signaling pathways and exhibit pronounced ligand-dependent biased agonism (Wang et al., 2018). The combination of large extracellular domains (ECD), extended orthosteric binding sites, and conformational flexibility makes class B1 GPCRs interesting but challenging targets for rational ligand discovery and particularly demanding benchmarks for computational ligand design and virtual screening methodologies (Aksu et al., 2024; M. Zhang et al., 2024).

Within this receptor family, the glucagon-like peptide-1 receptor (GLP-1R) plays a central role in the regulation of glucose homeostasis and energy metabolism. Glucagon-like peptide 1 (GLP-1) is a gut-derived peptide secreted from intestinal L-cells after a meal, which activates GLP-1R triggering multiple intracellular signaling cascades, predominantly through Gs-mediated cAMP signaling, thereby regulating insulin secretion, gastric emptying, appetite control, and carbohydrate metabolism (Drucker, 2018; Reed et al., 2021).

Several GLP-1R peptide-based agonists – including liraglutide, semaglutide, dulaglutide, and tirzepatide (a dual GLP-1R/GIPR agonist) – have been approved for the treatment of obesity and type 2 diabetes, achieving unprecedented improvements in glycemic control and body weight reduction (Arslanian et al., 2022; Horowitz et al., 2012; Jastreboff et al., 2022; Wilding et al., 2021). Despite their clinical efficacy, most GLP-1-derived peptides require parenteral administration (with oral semaglutide being the exception) and frequently induce gastrointestinal adverse events that limit long-term adherence (Aroda et al., 2019; Zhao et al., 2020). In addition, a recent meta-analysis has shown an overwhelming weight recovery after GLP1RA treatment. These limitations have motivated sustained efforts to develop orally available non-peptidic GLP-1R agonists.

Nevertheless, the discovery of small-molecule GLP-1R agonists has proven challenging, as such ligands must reproduce the functional consequences of large peptides that stabilize the active receptor through numerous and spatially distributed interactions. Structural studies have shown that GLP-1R activation involves a complex network of interactions spanning the ECD and the transmembrane (TM) core, including deep penetration of the GLP-1 N-terminus into the orthosteric cavity and stabilization of characteristic active-state rearrangements within the TM bundle (Cong et al., 2022; X. Zhang et al., 2020). This structural complexity, together with the availability of multiple active-state conformations, makes GLP-1R a stringent test case for assessing the robustness of computational screening strategies.

Early non-peptidic compounds such as TT-OAD2 provided initial proof that direct engagement of the TM pocket by small molecules could promote GLP-1R activation (Zhao et al., 2020). More advanced non-peptidic agonists have been identified with distinct binding modes and activation mechanisms. One notable example is orforglipron (LY3502970), a small molecule in Phase III trial, binds to GLP-1R at a distinct site closer to the extracellular side of the transmembrane helical bundle rather than deeply within the orthosteric pocket (Kawai et al., 2020; X. Zhang et al., 2020). Unlike peptide agonists such as liraglutide or semaglutide, orforglipron engages a different set of receptor residues and stabilizes an active receptor conformation associated with a signaling profile broadly similar to that of GLP-1, although arising from a distinct binding mode and receptor interaction network (Cong et al., 2022). Another similar non-peptidic candidate, danuglipron (PF-06882961), was rationally designed to occupy the deep orthosteric core of the receptor, overlapping with the binding region of the N-terminal residues of GLP-1 that penetrate into the transmembrane cavity. Structural analyses revealed that danuglipron stabilizes a receptor conformation distinct from those induced by either GLP-1 or orforglipron, illustrating the existence of multiple active-state geometries compatible with GLP-1 activation (Griffith et al., 2022). Although the development of danuglipron was discontinued in 2025 (Pfizer, 2025).

Identifying small-molecule agonists for class B1 GPCRs such as GLP-1R remains challenging due to the receptor’s conformational plasticity and its large, peptide-optimized binding interface (Liu et al., 2025). Therefore, despite major advances in computational drug discovery, conventional virtual screening (VS) approaches that rely on a single method, similarity criterion, or receptor conformation are therefore prone to bias when applied to such structurally complex targets (Chen et al., 2024). In particular, single-method workflows often favor specific chemotypes or receptor states, potentially limiting scaffold diversity and reducing enrichment efficiency. Here, we address these challenges by implementing an integrated virtual screening workflow that combines complementary ligand-based and structure-based approaches and exploits consensus across multiple screening methodologies and active GLP-1R conformations. By explicitly leveraging methodological complementarity and multi-conformation consensus, the proposed framework aims to reduce conformational and chemotype bias while improving hit prioritization robustness. Using this strategy, we were able to identify three chemically distinct GLP-1R agonist candidates from large compound libraries: GQB47810 (non-peptidic small molecule), DPDPE (an endorphin-derived peptide), and neuromedin C, demonstrating the ability of the pipeline to capture diverse activation strategies and efficiently guide experimental validation.

## 2. Material and Methods

### 2.1. Library generation

Chemical structures from Drugbank (DB) v5.1.4 (https://go.drugbank.com/), FooDB (FDB) pre-1.0 (https://foodb.ca/), Biosynth (version as of March 1, 2024) (https://www.biosynth.com/), MolPort Natural (version as of April 13, 2024) (https://www.molport.com/shop/natural-compound-database), and Enamine HLL (v2021-https://enamine.net/compound-libraries/diversity-libraries/hit-locator-library-460) were collected and prepared for use in both ligand-based and structure-based computational workflows, amounting to more than one million compounds. All molecules were standardized by converting structures to 3D format, correcting protonation states at physiological pH, removing duplicates and inorganic entries, and generating low-energy conformations. The resulting curated libraries provided a chemically diverse set of ligand candidates suitable for downstream pharmacophore modeling, similarity searches, and docking-based virtual screening.

Compound libraries were converted into the required formats using a set of cheminformatics tools. Conversion to mol2 format was performed using the molconvert utility from ChemAxon included in JChem v15 (https://www.chemaxon.com/products/instant-jchem-suite/instant-jchem/) enabling 3D structure generation with low energy conformer search (S{fine}), mmff94 (Halgren, 1996) force-field minimization, and very strict geometry optimization criteria (L3). For pdbqt preparation, the prepare_ligand4.py script from AutoDockTools 1.5.7 (Trott & Olson, 2010) was employed, retaining all atoms without cleanup (-U) while repairing hydrogens (-A) to ensure compatibility with docking calculations. The.drs conversion template included in MetaScreener (https://github.com/bio-hpc/metascreener) was used for conversion with Dragon v6 (https://www.talete.mi.it/products/products.htm) for its own format.

For LigandScout-based pharmacophore screening (Wolber & Langer, 2004) libraries were converted to ldb format using the idbgen utility, applying the icon best preset and more (600) conformers to ensure adequate sampling of ligand flexibility. Finally, libraries used in the FPscreener/ConFiLiS (https://github.com/Jnelen/ConFiLiS) workflow were converted using the convert_library.py script provided within the ConFiLiS package, generating the internal fingerprint-compatible format required.

### 2.2. Protein Preparation

The crystal structure of human GLP-1R in a complex with 2 non-peptide agonists, TTOAD2 (PDB ID: 6ORV)(Zhao et al., 2020), LY3502970 (PDB ID: 6XOX)(Kawai et al., 2020) were obtained from the Protein Data Bank (https://www.rcsb.org/) (Goodsell et al., 2020). Protein structures were prepared using Maestro Schrödinger Suite (Schrödinger Release 2025-3: Maestro; Schrödinger). The Protein Preparation Wizard was used to assign bond orders, add hydrogens, generate zero-order bonds to metals, and correct missing side chains or loops when required. All crystallographic water molecules beyond 5 Å of the binding site were removed. Prepared receptor models were then processed in System Builder to define the simulation environment, assign default physiological ionization conditions, and were then subjected to a final restrained minimization to remove steric clashes and ensure structural relaxation before virtual screening calculations.

### 2.2. Structure-Based Virtual Screening

#### 2.2.1. Targeted docking

Targeted docking was performed using two active-state: GLP-1R structure in complex with TT-OAD2 (PDB ID: 6ORV), and GLP-1R structure in complex with orforglipron (PDB ID: 6XOX), to evaluate the binding modes of candidate compounds within the orthosteric binding site. Docking simulations were carried out using Lead Finder (LF) (Stroganov et al., 2008) implemented within the MetaScreener virtual screening platform (https://github.com/bio-hpc/metascreener), which enables automated, large-scale docking and post-processing of results.

Compound libraries from DB, FDB, and Biosynth were docked against both receptor models. For each structure, the docking grid was centered on the ligand-binding pocket defined by the co-crystallized agonist present in the corresponding crystal structure. Standard LF scoring functions were used to generate and rank multiple binding poses for each ligand. To increase robustness and reduce conformation-specific bias, docking results were compared across the two GLP-1R structures for each compound library. A consensus selection strategy was applied, retaining only compounds that exhibited reproducible binding poses and consistent interaction patterns in both receptor conformations. These consensus docking hits were prioritized for further analysis and incorporated into a multimethod consensus framework together with the other SBVS results (Figure 1).

**Figure 1.**
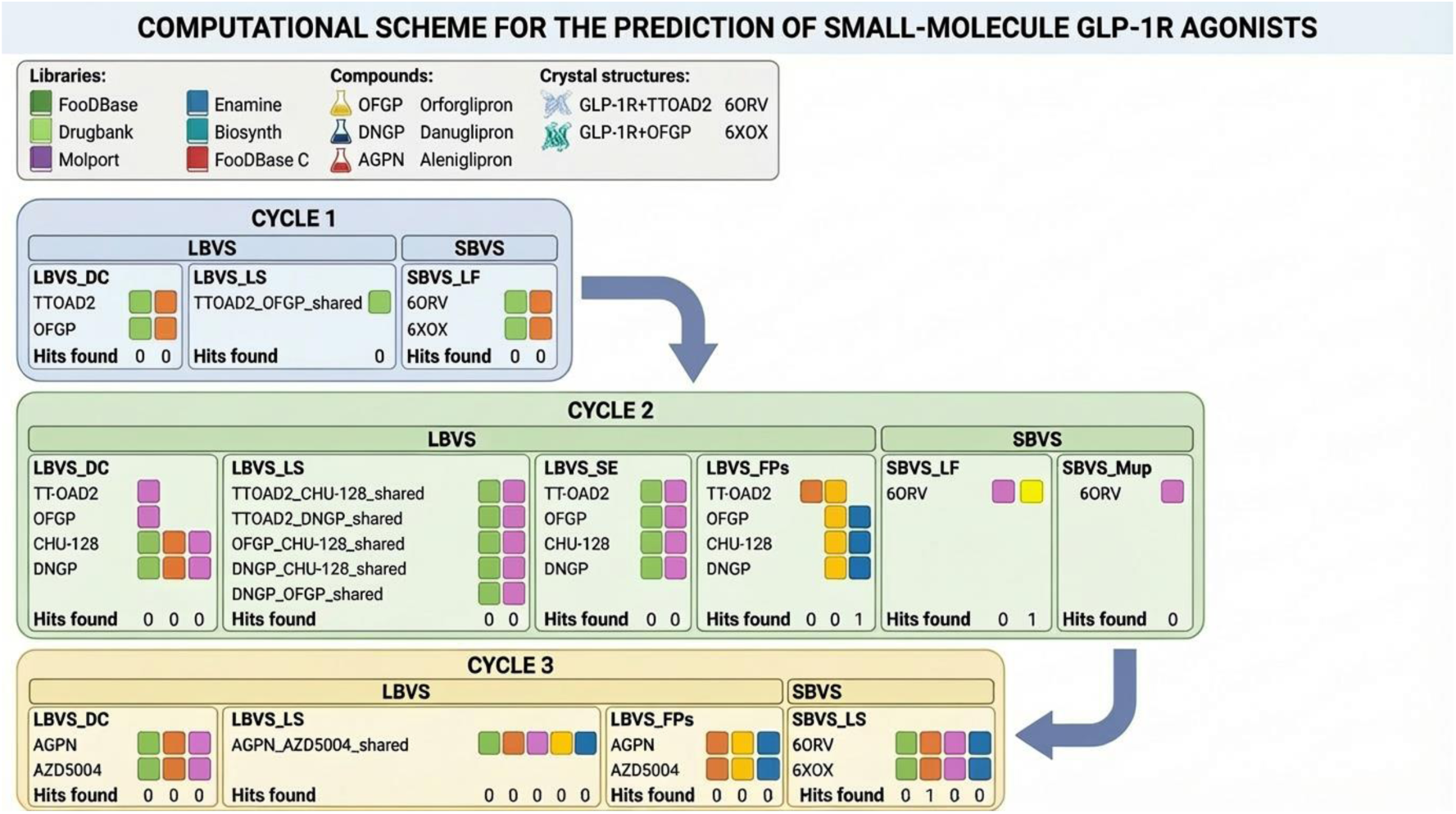
Computational scheme for the prediction of small-molecule GLP-1R agonists: Schematic overview of the integrated ligand-based (LBVS) and structure-based virtual screening (SBVS) workflow applied to identify candidate GLP-1R agonists. Compound libraries (FDB, DB, MolPort, Enamine, and Biosynth) were screened across three consecutive cycles using complementary ligand-based strategies, including descriptor-based methods (LBVS_DC), ligand similarity (LBVS_LS with shared pharmacological model), electrostatic similarity (LBVS_SE), and fingerprint-based screening (LBVS_FPs), as well as structure-based approaches such as molecular docking (SBVS_LF) and molecular pharmacophore modeling (SBVS_Mup). Active-state GLP-1R crystal structures in complex with reference agonists (PDB IDs 6ORV and 6XOX) were used for structure-based calculations. The number of prioritized candidates with proven in vitro activity (“hits found”) obtained at each step is indicated. The scheme illustrates the consensus-driven selection process that guided the identification of active compounds.

#### 2.2.2. Pharmacophore model generation

Structure-based pharmacophore models were generated using LigandScout (Wolber & Langer, 2004). The two active-state GLP-1R structures in complex with non-peptidic agonists (PDB IDs: 6ORV, 6XOX) were imported and processed to extract key ligand-receptor interaction features. For each complex, LigandScout automatically identified hydrogen-bond donors and acceptors, hydrophobic regions, aromatic interactions, and charged features based on the 3D arrangement of the bound peptide within the orthosteric binding pocket. A massive virtual screening calculation was performed using our in-house software MetaScreener (https://github.com/bio-hpc/metascreener), enabling automated, high-throughput screening of large compound libraries by parallelizing pharmacophore matching calculations across multiple ligands and receptor-derived models, thereby allowing efficient evaluation of thousands of candidate molecules. The pharmacophore models were screened against the DB, FDB, MolPort Natural and Biosynth libraries, and the maximum number of features to be omitted (-a,-*allow_omit*) for this calculation was 2. Our aim was to obtain compounds whose pharmacophore features matched the essential interaction pattern observed in the active-state GLP-1R structures. We applied a post-filtering step to obtain only those molecules with a pharmacophore score value greater than 0.70, selecting 200 compounds with the highest values. Finally, compounds were filtered based on commercial availability in the MolPort catalog. This filtering step yielded four compounds from DB, six from FDB, three from MolPort Natural, and eight from Biosynth.

### 2.3. Ligand-based virtual screening

Ligand-based virtual screening (LBVS) was performed to identify small-molecule candidates with physicochemical, structural, or pharmacophoric similarity to known GLP-1R agonists. A panel of reference ligands representing diverse chemotypes – including TT-OAD2, PF-06882961 (danuglipron), LY3502970 (orforglipron), CHU-128, GSBR-1290 (aleniglipron), and AZD5004 (Figure 2) – was used to derive the ligand-centered models described below. All calculations were performed using curated and standardized libraries from DB, FDB, Biosynth, Molport and Enamine.

**Figure 2.**
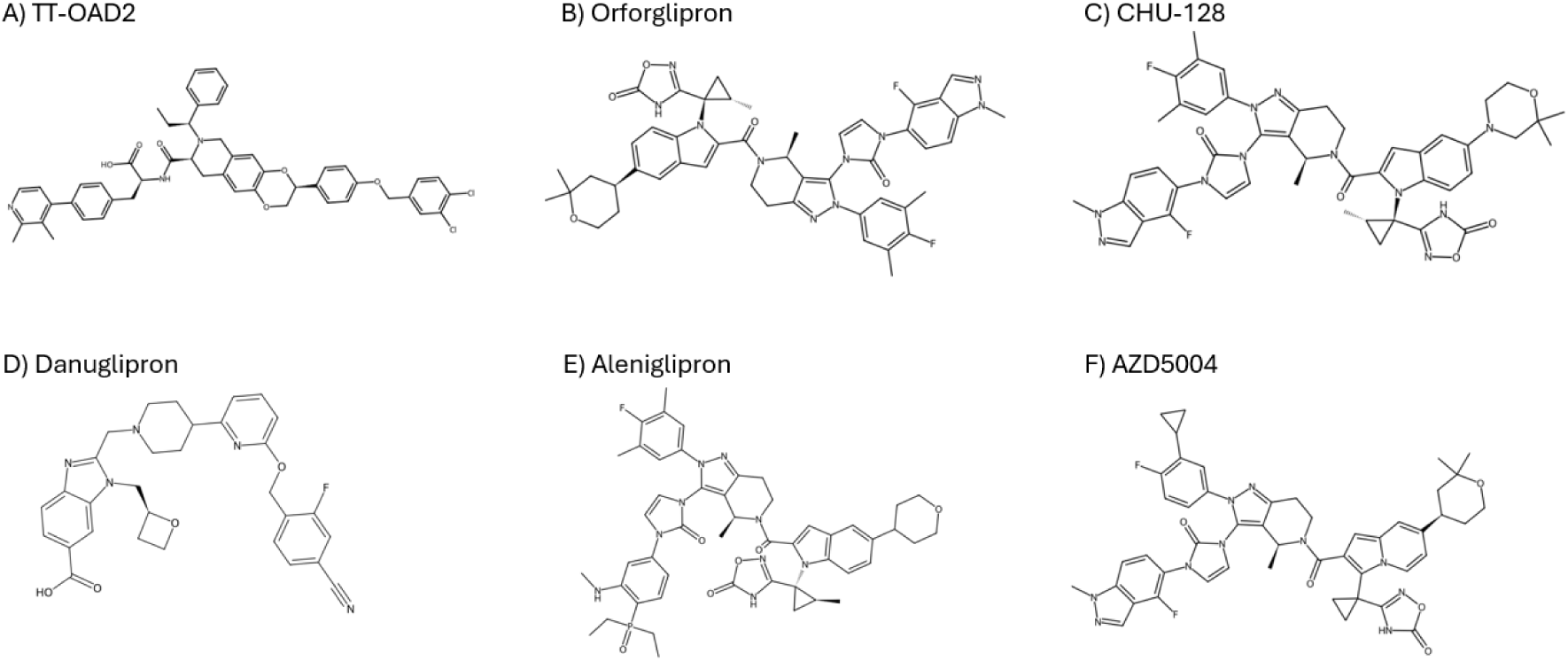
2D chemical structures of reference control compounds reported as nonpeptidic ligands of the GLP-1R. A) TT-OAD2; B) Orforglipron; C) CHU-128; D) Danuglipron; E) Aleniglipron; and F) AZD5004, which were used as positive controls for comparative analyses.

#### 2.3.1. Descriptor-based screening

A descriptor-based analysis was conducted to compare global physicochemical profiles between all reference agonists and candidate molecules from DB, FDB, and Molport libraries. For all compounds, molecular descriptors – including molecular weight, logP, topological polar surface area (tPSA), hydrogen bond donors/acceptors, rotatable bonds, aromaticity indices, and charge-related parameters – were computed using QikProp from Maestro Desmond (Schrödinger Release 2024-3: QikProp, 2025). Candidate molecules were ranked by their Euclidean distance to the descriptor centroid of reference ligands. Compounds falling within the predefined similarity threshold were retained for subsequent evaluations.

#### 2.3.2. Fingerprint similarity screening (FPscreener)

Two-dimensional structural similarity screening was performed using FPscreener (https://github.com/Jnelen/ConFiLiS), a tool enabling multi-fingerprint analysis (Nelen et al., 2025). Circular (ECFP-like), topological (FP2/FP4), and path-based fingerprints were generated for all reference ligands and Drugbank, Enamine and Biosynth libraries. Similarity was quantified using Tanimoto coefficients, and a Consensus_Average score – combining results from all fingerprint types – was used to prioritize hits. Compounds displaying consistent similarity across multiple fingerprint classes were selected for downstream filtering.

#### 2.3.3. Electrostatic similarity analysis

Electrostatic similarity calculations were performed using OpenEye toolkit by combining ROCS for shape-based alignment and EON for electrostatic similarity calculations. ROCS was first applied to align candidate molecules from FDB and Molport libraries to the reference GLP-1R agonists (TT-OAD2, LY3502970, PF-06882961 and CHU-128) and to rank them according to the TanimotoCombo score, which integrates shape and chemical feature overlap. The top 2000 ROCS-ranked hits were retained and exported in mol2 format for subsequent electrostatic evaluation.

Electrostatic similarity between each candidate and the corresponding reference ligand was then computed using EON based on molecular electrostatic potential fields. Compounds were ranked according to their Electrostatic Tanimoto (ET) similarity score. Molecules exhibiting an ET score ≥ 0.7 were considered to display high electrostatic similarity to the reference agonist and were retained as potential candidates for further consensus-based prioritization. Visualization and manual inspection of selected overlays and electrostatic maps were performed using VIDA 5.0.4.1. (OpenEye Scientific Software, 2025).

#### 2.3.4. Ligand-based pharmacophore modeling

Ligand-based pharmacophore models were generated using LigandScout from conformationally optimized structures of reference GLP-1R agonists. For each ligand, LigandScout initially identified all potential pharmacophoric features, including hydrogen bond donors/acceptors, aromatic interactions, hydrophobic regions, and charged or ionizable groups. Since individual reference molecules exhibited a large number of features – reflecting their distinct chemotypes – direct merging into a single multi-ligand pharmacophore was not feasible.

To address this, pairwise shared-feature pharmacophore models were generated. Each model was constructed from pairs of reference ligands, retaining only the pharmacophoric features that were conserved between both molecules. This strategy reduced feature redundancy and emphasized interaction motifs that were recurrent across different chemotypes. The resulting set of pairwise shared pharmacophores was then used to screen the DB, FDB, Molport and Enamine libraries using MetaScreener. LigandScout provided a pharmacophore fit score for each screened molecule. To ensure high-confidence hit selection, only compounds achieving a fit score ≥ 0.70 for each model were retained and integrated into the multi-method consensus framework used to consolidate LBVS predictions.

#### 2.3.5. Bibliography searches

Before experimental evaluation, an exhaustive bibliographic search was conducted to assess whether the computationally prioritized compounds had been previously reported or associated with obesity, metabolic disorders, or incretin-related pathways. This analysis was performed to contextualize the predicted candidates within the existing literature and to identify any prior pharmacological or biological evidence relevant to their potential metabolic effects. The literature assessment served as an additional filtering and validation step, ensuring that subsequent in vitro characterization focused on compounds with either novel or insufficiently explored links to metabolic regulation.

### 2.5. In vitro pharmacology

#### 2.5.1. Cell culture

HEK293T cells were cultured in DMEM supplemented with 10% FBS and maintained in a humidified chamber with 5% CO_2_ at 37 °C.

#### 2.5.2. cAMP accumulation assay

Peptide– or small molecule-stimulated cAMP accumulation was measured by a LANCE Ultra cAMP kit (Revvity). Briefly, take the cells out of liquid nitrogen (all cells used in the experiment are stable cell lines). Once the cell suspension is completely thawed, add Stimulation Buffer (Hanks’ balanced salt solution (HBSS) supplemented with 5 mM HEPES, 0.5 mM IBMX and 0.1% (w/v) BSA, pH 7.4) and centrifuge at 1000 rpm for 5 minutes. Discard the supernatant and resuspend the cells with stimulation buffer to a density of 0.6 million cells per mL, then add the cells to 384-well white plates (3,000 cells per well). Different concentrations of ligand in stimulation buffer were added and the stimulation lasted for 40 min at RT. The reaction was stopped by adding 5 μL Eu-cAMP tracer and 5 μL ULight-anti-cAMP. After 1 h incubation at RT, the plate was read by an Envision plate reader (PerkinElmer) to measure TR-FRET signals (excitation: 320 nm, emission: 615 nm and 665 nm).

#### 2.5.3. β-arrestin1/2 recruitment assay

HEK293T cells were seeded at a density of 0.3 million cells per mL into 6 cm dish. After incubation for 24 h to reach 80% confluence, the cells were transiently transfected with GLP-1R-Rluc8 and β-arrestin 1/2-Venus at a 1:3 mass ratio using lipofectamine LTX reagent (Invitrogen) and cultured for 24 h. Then, seed the cells into white 96-well plates coated with poly-D-lysine hydrobromide at a density of 50000 cells per well and cultured for another 24h. Thereafter, cells were washed once and incubated for 30 min at 37℃ with HBSS buffer (pH 7.4) supplemented with 0.1% BSA and 10 mM HEPES. Five micromolars of coelenterazine h (Yeasen Biotech) was then added and incubated for 5 min in the dark. A 1.5-min baseline of BRET measurement was taken before the addition of ligand and BRET signal was measured at 10-s intervals for further 9 min. After removing baseline and background readings by subtracting average values of the baseline measurement and average values of vehicle-treated samples, respectively, the AUC across the time-course response curve was determined. Concentration-response curves were plotted using the total area-under-the-curve during the time of measurement post ligand addition.

#### 2.5.4. Receptor selectivity assay

Receptor selectivity was evaluated by comparing the ability of compounds to activate related incretin receptors. CHO cells stably expressing human GLP-1R or GIPR were used, and receptor activation was assessed by measuring cAMP accumulation under identical experimental conditions.

Compounds were tested across the same concentration ranges at both receptors, and responses were normalized to the maximal effect elicited by the corresponding endogenous ligand. This analysis enabled direct comparison of functional activity and allowed the identification of compounds displaying dual GLP-1R-selective or dual GLP-1R/GIPR agonistic profiles.

### 2.6. Statistical analysis

All functional assay data were analyzed using GraphPad Prism and are reported as mean ± SEM from at least three independent experiments. Concentration-response curves were fitted by nonlinear regression using three-parameter logistic equations. Statistical significance was assessed using two-tailed Student’s t test or one-way ANOVA, as appropriate for the comparison, and p <0.05 was considered statistically significant.

## 3. Results and Discussion

### 3.1. Computational prediction of novel GLP1R agonists

An integrated virtual screening workflow combining ligand-based and structure-based approaches was implemented to predict novel small-molecule agonists of GLP-1R, as shown in Figure 1. The strategy was designed to exploit methodological complementarity and reduce model-specific bias through consensus-driven prioritization across multiple screening routes.

Across the full screening pipeline, a total of 58 compounds were prioritized for experimental evaluation based on computational performance and commercial availability. These compounds were subsequently subjected to in vitro assays to assess GLP-1R activation. Among them, three compounds demonstrated reproducible agonistic activity, thereby validating the predictive capacity of the integrated computational strategy. The corresponding hit compounds are summarized in Figure 3.

**Figure 3.**
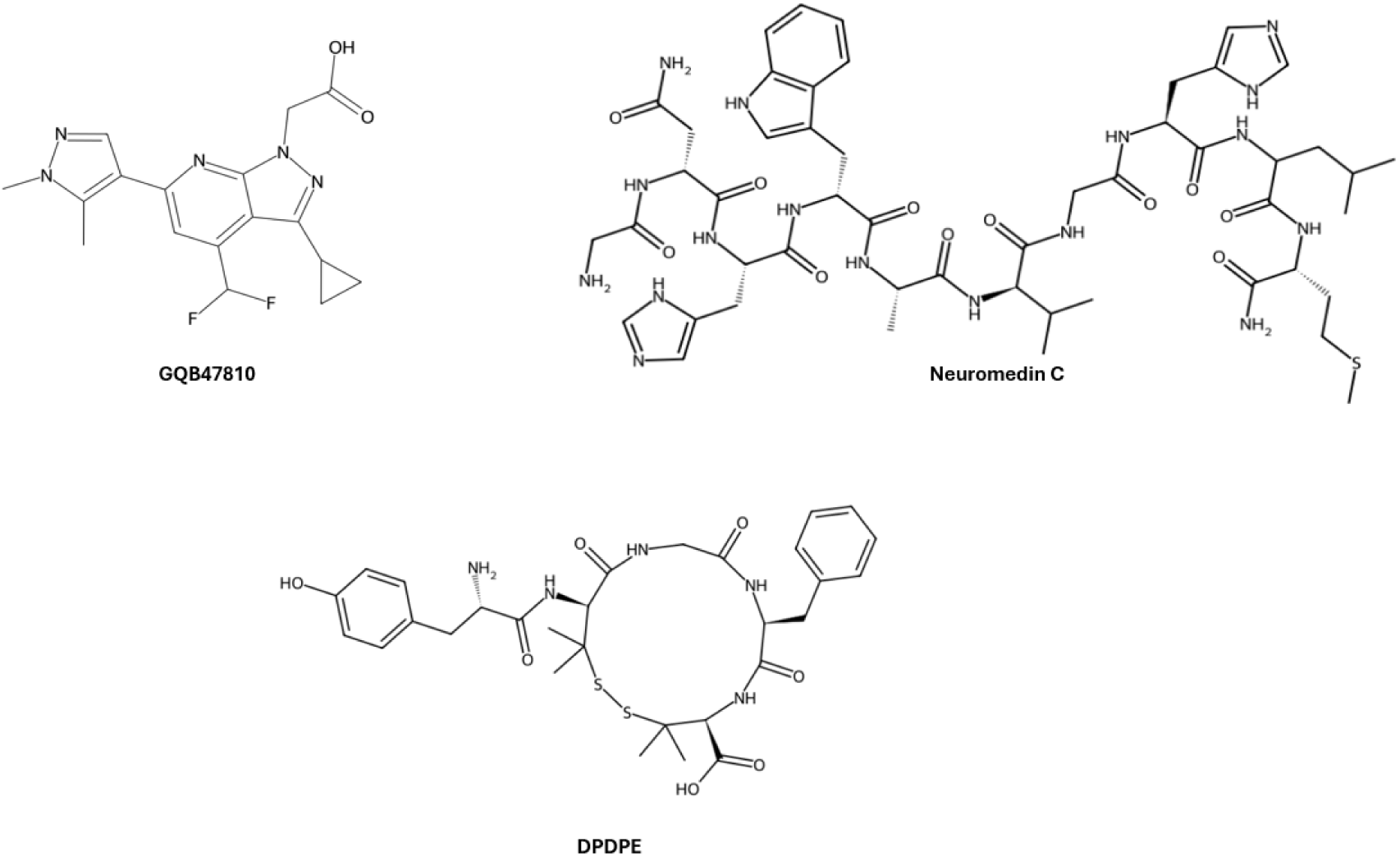
2D chemical structures of GLP-1R agonist candidates identified through complementary computational screening strategies. GQB47810, a LBVS hit predicted via multi-fingerprint consensus screening using FPscreener with orforglipron and CHU-128; neuromedin C, predicted through molecular docking; and DPDPE, a SBVS pharmacophore-derived hit predicted using LigandScout.

#### 3.1.1. Ligand-based prediction of GLP-1R agonist candidates

From the different consensus combinations evaluated, only the intersection of results obtained using LY3502970/CHU-128 consensus to Biosynth database resulted in the prioritization of a single ligand-based hit (GQB47810). The selected hit displayed a favorable multi-fingerprint similarity profile across Avalon, ECFP4, and PubChem fingerprints, as reflected in its overall average and consensus-normalized scores.

These results indicate that, while consensus screening was broadly applied, only specific combinations of reference ligands converged toward overlapping candidate sets, underscoring the importance of reference selection in ligand-based virtual screening and highlighting the complementarity of fingerprint-based similarity metrics for complex GPCR targets such as GLP-1R.

This represents a key aspect in the identification of novel compounds with GLP-1R agonistic activity, as those described to date exhibit markedly different receptor binding modes. Some compounds, such as danuglipron, display signaling profiles similar to the endogenous ligand, including B-arrestin recruitment and activation of additional downstream pathways (X. Zhang et al., 2020). In contrast, other molecules such as CHU-128, although binding in related receptor regions, are unable to fully recapitulate the pharmacological behavior of GLP-1. Furthermore, adding to the complexity of this process, allosteric activators have been described that bind to sites distinct from the orthosteric binding pocket of the receptor (Zhao et al., 2020), thereby necessitating a highly precise preselection of compounds to be used as lead candidates.

Following ligand-based prioritization, targeted molecular docking was performed to further characterize the predicted binding mode of GQB47810 within the GLP-1R orthosteric site. This analysis aimed to rationalize the selection of the compound by examining its ability to engage receptor residues previously implicated in GLP-1R activation and small-molecule agonist binding.

The predicted binding pose revealed that the compound establishes stabilizing interactions with a set of hydrophobic and aromatic residues lining the transmembrane binding cavity (Figure 4), including A200, L201, M204, L217, Y220, V229, and W297. These residues are located within the upper and central regions of the TM2 and TM3 bundle and have been recurrently associated with ligand recognition in active-state GLP-1R structures (Zhao et al., 2020). In particular, interactions with Y220 and W297, residues known to play a key role in stabilizing active receptor conformations, suggest that the compound is able to adopt a binding mode compatible with receptor activation.

**Figure 4.**
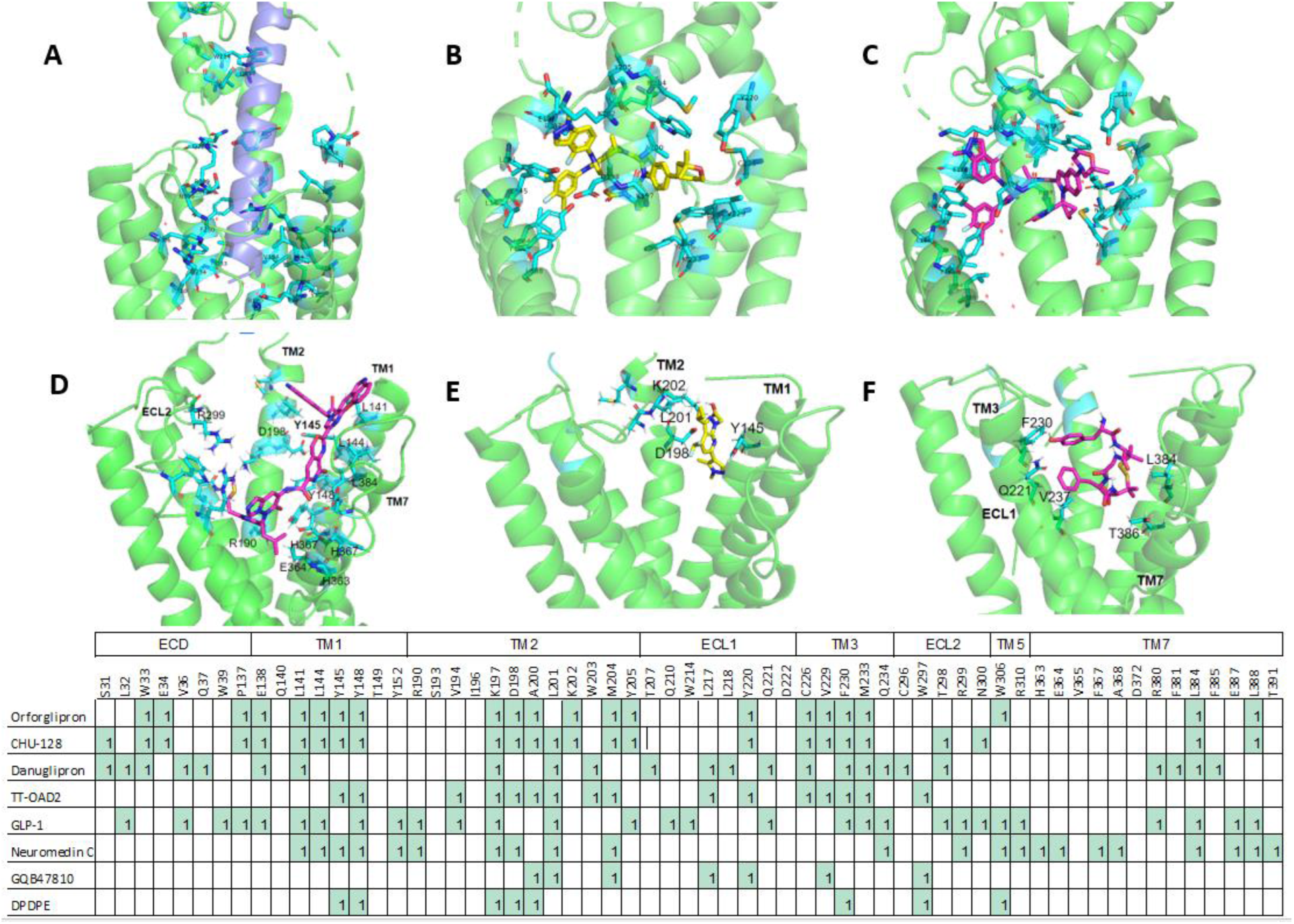
Structural interactions of GLP-1 receptor (GLP-1R) with peptide and small-molecule ligands. A) Binding interactions between GLP-1R and the endogenous peptide agonist GLP-1. The GLP-1 peptide is highlighted in purple, while interacting GLP-1R residues are shown in cyan. B)Interaction network between GLP-1R and the small-molecule agonist orforglipron. Orforglipron is shown in yellow and the interacting receptor residues in cyan. C) Binding interactions between GLP-1R and CHU-128. The ligand is represented in pink and interacting residues are shown in cyan. D) Interactions between GLP-1R and neuromedin C. Neuromedin C is depicted in pink and interacting receptor residues are shown in cyan. E) Interaction pattern between GLP-1R and compound GQB47810. The ligand is shown in yellow and receptor residues involved in the interaction are represented in cyan. F) Binding interactions between GLP-1R and DPDPE. DPDPE is represented in pink and the interacting GLP-1R residues are highlighted in cyan. The lower panel summarizes the interaction fingerprint between GLP-1R residues and the different ligands analyzed, indicating the residues involved in ligand recognition across receptor regions including the extracellular domain (ECD), transmembrane helices (TM), and extracellular loops (ECL)

The predominance of hydrophobic contacts observed in the predicted complex is consistent with binding modes reported for non-peptidic GLP-1R agonists, which typically engage the receptor through compact interaction networks within the transmembrane pocket rather than through extensive ECD contacts characteristic of peptide ligands. A more detailed comparison with structurally characterized non-peptidic GLP-1R agonists further contextualizes the predicted binding mode of GQB47810. In particular, the interaction pattern observed for this molecule shows notable similarities with LY3502970, which is reported to bind within the upper and central regions of the GLP-1R TM2 and TM3 bundle (Kawai et al., 2020). Structural studies of LY3502970 have highlighted the importance of hydrophobic and aromatic contacts within the cavity to stabilize active receptor conformations. Consistently, the predicted pose of the hit engages a comparable set of residues, suggesting partial overlap in receptor engagement despite differences in chemical scaffold.

Taken together, these observations indicate that GQB47810 preferentially targets an activation-relevant region of the GLP-1R TM cavity similar to that exploited by LY3502970, while remaining distinct from alternative binding modes exemplified by CHU-128. This structural convergence with known efficacious non-peptidic agonists provides additional support for the plausibility of the predicted binding mode and reinforces the relevance of the consensus-driven virtual screening strategy employed in this study. However, while the potency of GQB47810 strongly limits its therapeutic use (Figure 5), it paved the way for the development of new GLP1R agonists.

**Figure 5.**
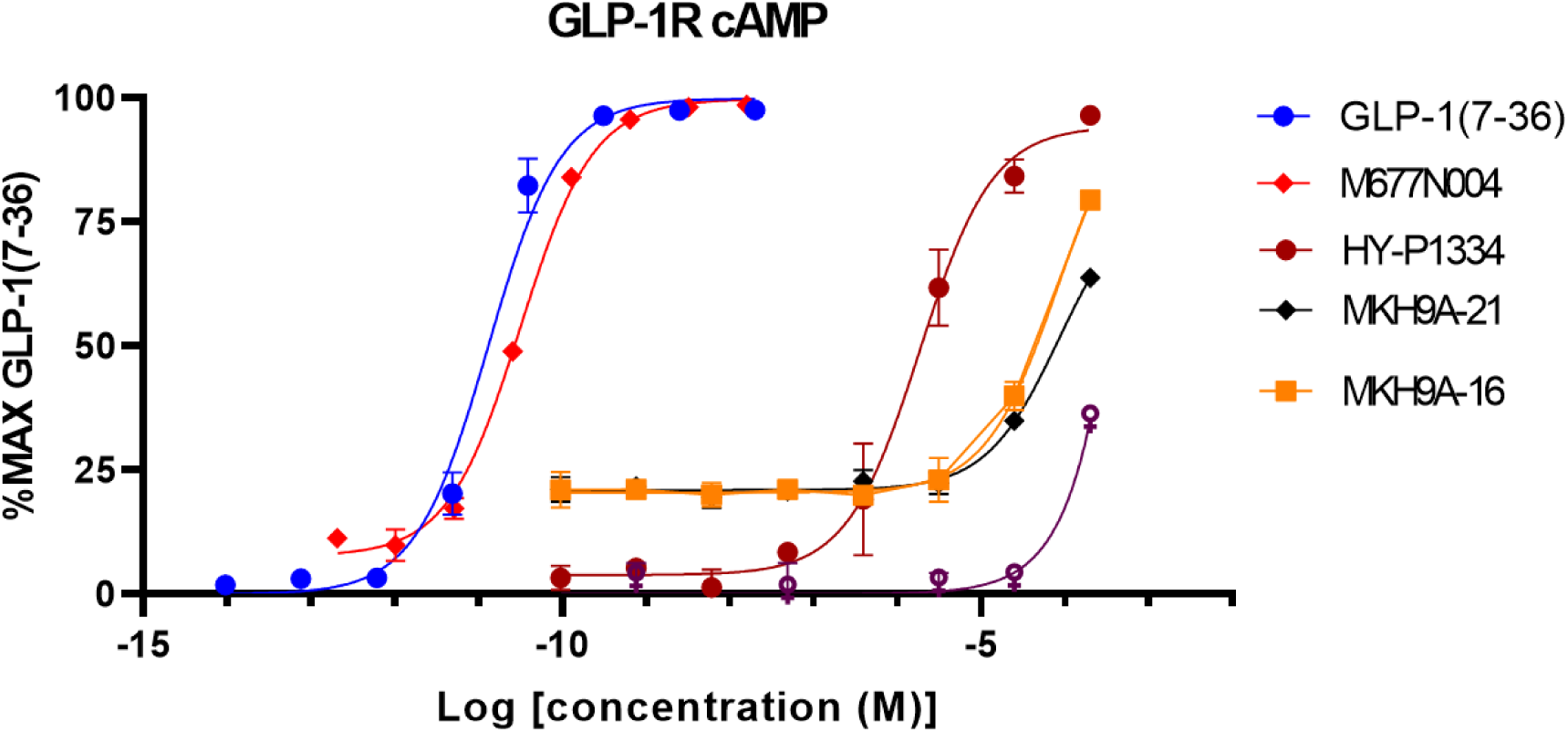
Dose-response curves showing the activation of GLP1R after stimulation by the endogenous ligand GLP1(7-36), a positive control M677N004 (danuglipron), HY-P1334 (DPDPE), MKH9A-21 (GQB47810), and MKH9A-16 (neuromedin C). Data show mean ± s.e.m of at least three replicates.

#### 3.1.2. Structure-based prediction identified two peptides as GLP1R agonists

SBVS of FDB using targeted molecular docking against the active-state GLP-1R structure 6ORV yielded a subset of five candidate compounds that consistently ranked among the top-scoring ligands. The inclusion of known control agonists within the screening set served to verify that the docking setup was able to recover ligands with established GLP-1R activity, thereby supporting the reliability of the prioritization strategy.

From these five compounds, neuromedin C was identified as a reproducible GLP-1R agonist, although, as happened with GQB47810, the in vitro data revealed a very low potency (Figure 5). The predicted binding mode places neuromedin C along the canonical peptide-binding groove, consistent with the class B1 GPCR “two-domain” recognition paradigm in which peptide ligands are initially oriented by extracellular receptor elements and subsequently stabilize the active TM architecture required for signaling (Figure 4). At the extracellular region, neuromedin C engages a cluster of hydrophobic and aromatic residues (L141, L144, Y145, Y148, Y152) consistent with peptide recognition and anchoring at the receptor surface. Additional contacts at the ECD-TM interface (R190, D198) suggest electrostatic stabilization at a coupling hotspot that can facilitate transmission of ligand binding into TM rearrangements. Within the TM 7 cavity, neuromedin C establishes interactions with residues lining the upper/mid helical bundle. The predicted interaction footprint extends further toward the intracellular region, yielding a broad contact network that plausibly stabilizes an activation-compatible receptor architecture from the extracellular face toward the cytoplasmic side (Figure 4).

This interaction pattern is consistent with endogenous-ligand agonism rather than typical non-peptidic binding modes (X. Zhang et al., 2020). Experimentally resolved GLP-1R complexes with peptide agonists (e.g., GLP-1, exendin-4, and clinically used GLP-1 analogs) commonly exhibit contacts distributed across extracellular regions and the TM bundle similar to those found with neuromedin C, being characterized by cooperative interactions that stabilize active conformations required for G protein coupling (Kawai et al., 2020).

The interaction mode of neuromedin C represent alternative activation strategies to the non-peptidic control agonists used in this study (TT-OAD2, danuglipron, LY3502970, and CHU-128), being characterized by a more compact and TM7-localized interaction networks. Relative to these non-peptidic controls, the docked neuromedin C displays a markedly broader footprint, including extracellular involvement, which is a hallmark of peptide-mediated receptor activation. However, this stronger association does not mean higher potency, as revealed by further in vitro assays. Moreover, the lack of interaction with key residues, such as W297 and R380 involved in agonist potency hampers the global effectiveness of neuromedin C as a full GLP1R agonist. Taken together, docking analyses suggest that neuromedin C engages GLP-1R through an interaction architecture resembling that of GLP1 and other peptide agonists, while remaining fundamentally distinct from the compact TM-centered binding patterns typical of non-peptidic ligands.

The compound that yielded the highest in vitro efficacy and potency was 2,5Pen-enkephalin (DPDPE) (Figure 5). This pentapeptide is analogous to enkephalin, an endogenous opioid that functions as neurotransmitter and neuromodulator in both the central and peripheral nervous systems to regulate different physiological processes like pain, mood, and reward behaviors (Stefanucci et al., 2017). According to the data derived from the SBVS, the interaction between DPDPE and GLP1R was characterized by a similar localization than danuglipron and orforglipron, within the TM bundle, and more concretely, closer to TM1-2 region (Figure 4). One unexpected data was the high efficacy of this compound, reaching similar levels to those observed with the endogenous ligand and danuglipron (Figure 5). However, further in vitro assays showed a different pharmacological profile, since although the effect on cAMP was notable, we did not observe additional significant effects on β-arrestin recruitment activation, therefore providing a similar profile than other non-peptidic agonists like CHU-128 and TT-OAD2. Again, several significant interactions were observed, like the hydrophobic interactions with TM1 residues Y145 and Y148, similar to TT-OAD2 and other peptidomimetic drugs, like Boc5 and WB4-24 (Zhao et al., 2020). However, other relevant interactions, especially those regulating the pharmacological potency of DPDPE, such as those with R380, a key residue for peptide affinity and receptor activation at TM7, was absent (Griffith et al., 2022). Interestingly, this residue is conserved in Class B1 GPCRs, such as glucagon, vasoactive intestinal peptide (VIP), and glucagon-like peptide-1 (GLP-1) (Liu et al., 2025).

### 3.2. In vitro results

#### 3.2.1 GLP-1 receptor activation

Several analyses were carried out to evaluate the functional activity of the computationally prioritized compounds at GLP-1R. From a batch of 58 compounds, three showed reproducible, concentration-dependent stimulation of receptor signaling (Figure 5).

Among the activated compounds, only DPDPE displayed a clear concentration-dependent activation of human GLP-1R (Figure 5). The resulting concentration-response curve exhibited a characteristic sigmoidal profile, reaching maximal efficacy comparable to that of the endogenous ligand GLP-1(7-36). The compound achieved nearly 100% of the GLP-1 maximal response, indicating full agonistic behavior in terms of efficacy; Relative to GLP-1 and danuglipron, the curve exhibited a pronounced rightward shift, reflecting a substantial decrease in potency; specifically, the EC50 was in the 1-μM range. Despite this reduced potency, DPDPE consistently produced robust maximal activation, supporting its ability to stabilize and active receptor conformation. The preservation of full efficacy, despite lower potency, suggests that the compound effectively engages key activation-relevant residues within the receptor binding cavity, in agreement with predicted computational analyses.

In contrast to the lead candidate described above, GQB47810 and neuromedin C, exhibited lower GLP-1R agonistic activity (Figure cAMP). Both compounds produced only modest, concentration-dependent increases in cAMP accumulation and failed to reach maximal responses comparable to GLP-1.

#### 3.2.2. Receptor Selectivity Assays

As DPDPE showed the highest efficacy results, further in vitro studies were carried out to evaluate potential activities on other incretin-related receptors, as occur with dual agonist like tirzepatide (GLP1R/GIPR), survodutide (GLP1R/GCGR) and other experimental agonists like THDBH120 (GLP1R/GIPR) and MEDI0382 and SAR425899 (GLP1R/GCGR) (Zikhai Zheng et al., 2024; Yang Li et al., 2023; Feng Zhang et al., 2026; Jonathan E Campbell et al., 2023). In this regard, our data showed a similar pharmacological profile on glucose-dependent insulinotropic polypeptide receptor (GIPR), with similar efficacy than with GLP-1R, indicating a dual agonist profile rather than strict GLP-1R selectivity, as shown in Figure 6. Both the effectiveness and potency of GIPR was similar to that observed in the GLP1R. This balanced activity suggests that DPDPE is capable of engaging conserved activation mechanisms shared between incretin receptors, despite differences in their primary sequence and ligand recognition features (Figure Xa). Further analyses were conducted on glucagon receptor (GCGR), but at this time, DPDPE failed to show any activity on this receptor (data not shown).

**Figure 6.**
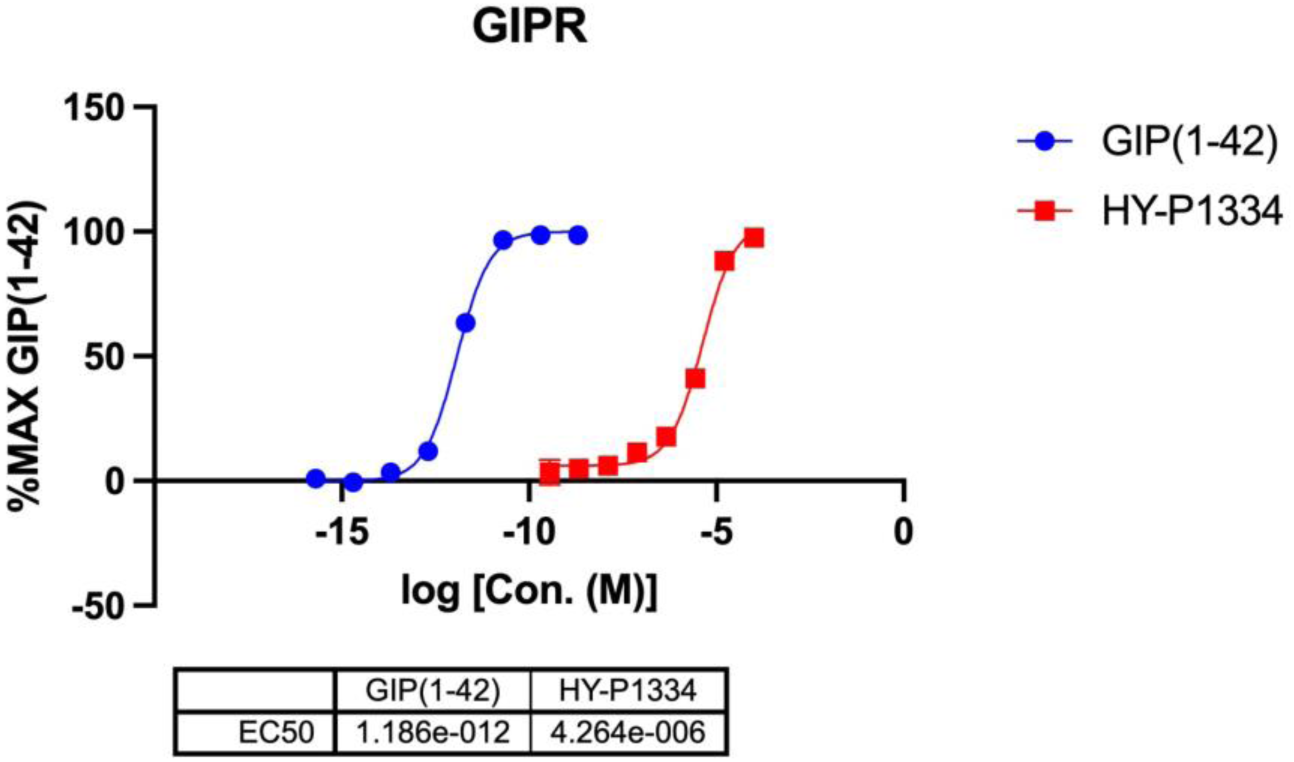
Concentration-response curves of DPDPE (HY-P1334) at human GLP-1R. Gs-mediated cAMP accumulation is expressed as the percentage of the maximal response elicited by GLP-1(7–36). GLP-1 (blue circles) displays high potency and full efficacy, whereas the lead compound (red squares) achieves comparable maximal efficacy with a rightward shift in the curve, indicating reduced potency. Data are presented as mean ± SEM from independent experiments.

The observed GLP-1R/GIPR dual activity is particularly similar to that of tirzepatide, a clinically approved dual incretin agonist (Jastreboff A et al., 2022). Importantly, tirzepatide is a large peptide-based agonist that engages both receptors through extensive and distributed interactions across ECD and transmembrane core (Coskun et al., 2022). In contrast, DPDPE is a small peptide, implying a fundamentally different mode of receptor engagement that likely involves a more localized interaction network within the transmembrane binding region. Therefore, it can be speculated that DPDPE or DPDPE-derivatives with higher potency may be more effective than current single-receptor agonists, like semaglutide.

We report here the first empirical evidence of a novel small peptide that functions as a bi-specific agonist for both GLP-1R and GIPR in vitro. The ability of a small peptide to reproduce aspects of dual incretin receptor activation in vitro, underscores its potential relevance as a lead scaffold. These findings further support the value of computationally guided discovery strategies for identifying ligands with complex and therapeutically relevant signaling profiles at class B1 GPCRS.

Furthermore, the application of a multifunctional agonist targeting the GLP-1, GIP, and delta-opioid (DOR) receptors—such as DPDPE—represents a novel therapeutic strategy for obesity, particularly given that DOR activation has been shown to independently improve glucose homeostasis (Olianas et al., 2011) and insulin sensitivity (Meulebrouck et al., 2024), as well as regulating emotional aspects of eating behavior (Wassum et al., 2009; Yoshioka et al., 2024). However, the high potency on DOR compared to the observed on GLP1/GIP receptors suggests that effective doses may saturate opioid receptors (Suh et al. described an EC50 of DPDPE on DOR = 5.2 ± 1.2 nM) (Suh & Tseng, 1990). However, these aspects will need to be explored in further studies to identify the most suitable doses to achieve the greatest possible effect on appetite regulation, inhibiting both hedonic and physiological cues, with the least possible sedative effect.

## 4. Conclusions

In this study, we present a robust and integrative virtual screening strategy for the identification of agonists of the GLP-1R. By combining complementary LBVS and SBVS approaches and explicitly exploiting consensus across multiple methods and receptor conformations, the proposed workflow addresses key limitations of conventional single-method screening strategies when applied to structurally complex class B1 GPCRs. This multi-layered approach allowed the efficient prioritization of three candidates from large chemical libraries, supporting its applicability as a general framework for early-stage GLP-1R ligand discovery.

From all the potential hits, DPDPE emerged as the lead candidate, achieving full agonism at the GLP-1R with a maximal response (Emax) equivalent to GLP-1 and danuglipron. Despite a reduction in potency relative to these standards, the observed GIPR activity suggests its potential as a molecular starting point for the rational design of triple-acting ligands (GLP-1R/GIPR/DOR). This integrated pharmacological profile aims to modulate both homeostatic and hedonic pathways of food intake, offering a first-in-class therapeutic paradigm for obesity management, as well as other appetite-related disorders.

Notably, this study establishes a new step in GLP-1R ligand design by demonstrating that small-peptide architectures can retain full functional efficacy, provided key pharmacophoric motifs are preserved. These findings challenge the prevailing paradigm that bulky peptide scaffolds are indispensable for receptor activation, significantly expanding the chemical space toward non-peptidic ligands with superior developability profiles.

Overall, the computational framework presented here offers a robust and generalizable blueprint for the rational discovery of incretin mimetics. Beyond the specific leads identified, this work underscores the power of consensus-based virtual screening to uncover non-intuitive solutions to complex molecular recognition challenges, effectively streamlining the development of next-generation GLP-1R modulators.

## Author contribution

E.M.G: Conceptualization. Formal analysis. Investigation. Methodology. Data curation. Writing – original draft, writing – review and editing; N.T: Investigation. Methodology. Writing – review and editing; J.R.A: Formal analysis. Methodology. Writing – review and editing; X.C: Investigation. Methodology. Resources. Writing – review and editing; D.Y: Investigation. Methodology. Resources. Validation. Writing – review and editing; J.J.H.M: Conceptualization. Investigation. Supervision. Validation. Writing – review and editing; H.P.S: Funding acquisition. Resources. Supervision. Validation. Writing – review and editing.

## Acknowledgements

The supercomputing resources used in this work have been supported by the Plataforma Andaluza de Bioinformatica of the University of Malaga, by the supercomputing infrastructure of the NLHPC (ECM-02, Powered@NLHPC), and by BSC.

## References

1. Aksu, H., Demirbilek, A., & Uba, A. I. (2024). Insights into the structure and activation mechanism of some class B1 GPCR family members. Molecular Biology Reports, 51(1), 966-. 10.1007/s11033-024-09876-w

2. Aroda, V. R., Rosenstock, J., Terauchi, Y., Altuntas, Y., Lalic, N. M., Villegas, E. C. M., Jeppesen, O. K., Christiansen, E., Hertz, C. L., & Haluzík, M. (2019). PIONEER 1: Randomized Clinical Trial of the Efficacy and Safety of Oral Semaglutide Monotherapy in Comparison With Placebo in Patients With Type 2 Diabetes. Diabetes Care, 42(9), 1724–1732. 10.2337/DC19-0749

3. Arslanian, S. A., Hannon, T., Zeitler, P., Chao, L. C., Boucher-Berry, C., Barrientos-Pérez, M., Bismuth, E., Dib, S., Cho, J. I., & Cox, D. (2022). Once-Weekly Dulaglutide for the Treatment of Youths with Type 2 Diabetes. The New England Journal of Medicine, 387(5), 433–443. 10.1056/NEJMOA2204601

4. Bortolato, A., Doré, A. S., Hollenstein, K., Tehan, B. G., Mason, J. S., & Marshall, F. H. (2014). Structure of Class B GPCRs: new horizons for drug discovery. British Journal of Pharmacology, 171(13), 3132–3145. 10.1111/BPH.12689

5. Chen, Z., Ren, X., Zhou, Y., & Huang, N. (2024). Exploring structure-based drug discovery of GPCRs beyond the orthosteric binding site. HLife, 2(5), 211–226. 10.1016/J.HLIFE.2024.01.002

6. Cong, Z., Zhou, Q., Li, Y., Chen, L. N., Zhang, Z. C., Liang, A., Liu, Q., Wu, X., Dai, A., Xia, T., Wu, W., Zhang, Y., Yang, D., & Wang, M. W. (2022). Structural basis of peptidomimetic agonism revealed by small-molecule GLP-1R agonists Boc5 and WB4-24. Proceedings of the National Academy of Sciences of the United States of America, 119(20), e2200155119. 10.1073/pnas.2200155119

7. de Graaf, C., Donnelly, D., Wootten, D., Lau, J., Sexton, P. M., Miller, L. J., Ahn, J. M., Liao, J., Fletcher, M. M., Yang, D., Brown, A. J. H., Zhou, C., Deng, J., & Wang, M. W. (2016). Glucagon-like peptide-1 and its class B G protein-coupled receptors: A long march to therapeutic successes. Pharmacological Reviews, 68(4), 954–1013. 10.1124/pr.115.011395

8. Drucker, D. J. (2018). Mechanisms of Action and Therapeutic Application of Glucagon-like Peptide-1. Cell Metabolism, 27(4), 740–756. 10.1016/j.cmet.2018.03.001

9. Goodsell, D. S., Zardecki, C., Di Costanzo, L., Duarte, J. M., Hudson, B. P., Persikova, I., Segura, J., Shao, C., Voigt, M., Westbrook, J. D., Young, J. Y., & Burley, S. K. (2020). RCSB Protein Data Bank: Enabling biomedical research and drug discovery. Protein Science: A Publication of the Protein Society, 29(1), 52–65. 10.1002/PRO.3730

10. Griffith, D. A., Edmonds, D. J., Fortin, J. P., Kalgutkar, A. S., Kuzmiski, J. B., Loria, P. M., Saxena, A. R., Bagley, S. W., Buckeridge, C., Curto, J. M., Derksen, D. R., Dias, J. M., Griffor, M. C., Han, S., Jackson, V. M., Landis, M. S., Lettiere, D., Limberakis, C., Liu, Y., … Tess, D. A. (2022). A Small-Molecule Oral Agonist of the Human Glucagon-like Peptide-1 Receptor. Journal of Medicinal Chemistry, 65(12), 8208–8226. 10.1021/ACS.JMEDCHEM.1C01856/SUPPL_FILE/JM1C01856_S I_003.CSV

11. Halgren, T. A. (1996). Performance of MMFF94*. Scope, Parameterization, and Journal of Computational Chemistry, 17, 490–519. 10.1002/(SICI)1096-987X(199604)17:5/6

12. Horowitz, M., Flint, A., Jones, K. L., Hindsberger, C., Rasmussen, M. F., Kapitza, C., Doran, S., Jax, T., Zdravkovic, M., & Chapman, I. M. (2012). Effect of the once-daily human GLP-1 analogue liraglutide on appetite, energy intake, energy expenditure and gastric emptying in type 2 diabetes. Diabetes Research and Clinical Practice, 97(2), 258–266. 10.1016/J.DIABRES.2012.02.016

13. Jastreboff, A. M., Aronne, L. J., Ahmad, N. N., Wharton, S., Connery, L., Alves, B., Kiyosue, A., Zhang, S., Liu, B., Bunck, M. C., & Stefanski, A. (2022). Tirzepatide Once Weekly for the Treatment of Obesity. The New England Journal of Medicine, 387(3), 205–216. 10.1056/NEJMOA2206038

14. Kawai, T., Sun, B., Yoshino, H., Feng, D., Suzuki, Y., Fukazawa, M., Nagao, S., Wainscott, D. B., Showalter, A. D., Droz, B. A., Kobilka, T. S., Coghlan, M. P., Willard, F. S., Kawabe, Y., Kobilka, B. K., & Sloop, K. W. (2020). Structural basis for GLP-1 receptor activation by LY3502970, an orally active nonpeptide agonist. Proceedings of the National Academy of Sciences of the United States of America, 117(47), 29959–29967. 10.1073/pnas.2014879117

15. Liu, T., Naidoo, N. R., Agyemang, E., & Lamichhane, R. (2025). Insights into the structural dynamics of the secretin family (class B1) G protein-coupled receptors. Journal of Biological Chemistry, 301(8), 110466. 10.1016/J.JBC.2025.110466

16. Meulebrouck, S., Merrheim, J., Queniat, G., Bourouh, C., Derhourhi, M., Boissel, M., Yi, X., Badreddine, A., Boutry, R., Leloire, A., Toussaint, B., Amanzougarene, S., Vaillant, E., Durand, E., Loiselle, H., Huyvaert, M., Dechaume, A., Scherrer, V., Marchetti, P., … Bonnefond, A. (2024). Functional genetics reveals the contribution of delta opioid receptor to type 2 diabetes and beta-cell function. Nature Communications 2024 15:1, 15(1), 6627-. 10.1038/s41467-024-51004-6

17. Nelen, J., Naponelli, V., Villalgordo-Soto, J. M., Falasca, M., & Pérez-Sánchez, H. (2025). Targeting Drug Resistance in Cancer: Dimethoxycurcumin as a Functional Antioxidant Targeting ABCC3. *Antioxidants (Basel*, Switzerland), 14(5). 10.3390/ANTIOX14050599

18. Olianas, M. C., Dedoni, S., & Onali, P. (2011). δ-Opioid receptors stimulate GLUT1-mediated glucose uptake through Src– and IGF-1 receptor-dependent activation of PI3-kinase signalling in CHO cells. British Journal of Pharmacology, 163(3), 624–637. 10.1111/J.1476-5381.2011.01234.X

19. Reed, J., Bain, S., & Kanamarlapudi, V. (2021). A Review of Current Trends with Type 2 Diabetes Epidemiology, Aetiology, Pathogenesis, Treatments and Future Perspectives. *Diabetes*, Metabolic Syndrome and Obesity: Targets and Therapy, 14, 3567–3602. 10.2147/DMSO.S319895

20. Stefanucci, A., Novellino, E., Mirzaie, S., Macedonio, G., Pieretti, S., Minosi, P., Szűcs, E., Erdei, A. I., Zádor, F., Benyhe, S., & Mollica, A. (2017). Opioid Receptor Activity and Analgesic Potency of DPDPE Peptide Analogues Containing a Xylene Bridge. ACS Medicinal Chemistry Letters, 8(4), 449. 10.1021/acsmedchemlett.7b00044

21. Stroganov, O. V., Novikov, F. N., Stroylov, V. S., Kulkov, V., & Chilov, G. G. (2008). Lead Finder: An Approach To Improve Accuracy of Protein−Ligand Docking, Binding Energy Estimation, and Virtual Screening. Journal of Chemical Information and Modeling, 48(12), 2371–2385. 10.1021/CI800166P

22. Suh, H. H., & Tseng, L. F. (1990). Different types of opioid receptors mediating analgesia induced by morphine, DAMGO, DPDPE, DADLE and β-endorphin in mice. Naunyn-Schmiedeberg’s Archives of Pharmacology, 342(1), 67–71. 10.1007/BF00178974

23. Trott, O., & Olson, A. J. (2010). AutoDock Vina: improving the speed and accuracy of docking with a new scoring function, efficient optimization and multithreading. Journal of Computational Chemistry, 31(2), 455. 10.1002/JCC.21334

24. Wang, W., Qiao, Y., & Li, Z. (2018). New Insights into Modes of GPCR Activation. Trends in Pharmacological Sciences, 39(4), 367–386. 10.1016/j.tips.2018.01.001

25. Wassum, K. M., Ostlund, S. B., Maidment, N. T., & Balleine, B. W. (2009). Distinct opioid circuits determine the palatability and the desirability of rewarding events. Proceedings of the National Academy of Sciences of the United States of America, 106(30), 12512–12517. 10.1073/PNAS.0905874106;WEBSITE:WEBSITE:PNAS-SITE;WGROUP:STRING:PUBLICATION

26. Wilding, J. P. H., Batterham, R. L., Calanna, S., Davies, M., Van Gaal, L. F., Lingvay, I., McGowan, B. M., Rosenstock, J., Tran, M. T. D., Wadden, T. A., Wharton, S., Yokote, K., Zeuthen, N., & Kushner, R. F. (2021). Once-Weekly Semaglutide in Adults with Overweight or Obesity. The New England Journal of Medicine, 384(11), 989–1002. 10.1056/NEJMOA2032183

27. Wolber, G., & Langer, T. (2004). LigandScout: 3-D Pharmacophores Derived from Protein-Bound Ligands and Their Use as Virtual Screening Filters. Journal of Chemical Information and Modeling, 45(1), 160–169. 10.1021/CI049885E

28. Yoshioka, T., Yamada, D., Hagiwara, A., Kajino, K., Iio, K., Saitoh, T., Nagase, H., & Saitoh, A. (2024). Delta opioid receptor agonists activate PI3K–mTORC1 signaling in parvalbumin-positive interneurons in mouse infralimbic prefrontal cortex to exert acute antidepressant-like effects. Molecular Psychiatry 2024 30:5, 30(5), 2038–2048. 10.1038/s41380-024-02814-z

29. Zhang, M., Chen, T., Lu, X., Lan, X., Chen, Z., & Lu, S. (2024). G protein-coupled receptors (GPCRs): advances in structures, mechanisms and drug discovery. Signal Transduction and Targeted Therapy 2024 9:1, 9(1), 88-. 10.1038/s41392-024-01803-6

30. Zhang, X., Belousoff, M. J., Zhao, P., Kooistra, A. J., Truong, T. T., Ang, S. Y., Underwood, C. R., Egebjerg, T., Šenel, P., Stewart, G. D., Liang, Y. L., Glukhova, A., Venugopal, H., Christopoulos, A., Furness, S. G. B., Miller, L. J., Reedtz-Runge, S., Langmead, C. J., Gloriam, D. E., … Wootten, D. (2020). Differential GLP-1R Binding and Activation by Peptide and Non-peptide Agonists. Molecular Cell, 80(3), 485–500.e7. 10.1016/j.molcel.2020.09.020

31. Zhao, P., Liang, Y. L., Belousoff, M. J., Deganutti, G., Fletcher, M. M., Willard, F. S., Bell, M. G., Christe, M. E., Sloop, K. W., Inoue, A., Truong, T. T., Clydesdale, L., Furness, S. G. B., Christopoulos, A., Wang, M. W., Miller, L. J., Reynolds, C. A., Danev, R., Sexton, P. M., & Wootten, D. (2020). Activation of the GLP-1 receptor by a non-peptidic agonist. Nature 2020 577:7790, 577(7790), 432–436. 10.1038/s41586-019-1902-z

